# Matching sounds to shapes: Evidence of the Bouba-Kiki effect in naïve baby chicks

**DOI:** 10.1101/2024.05.17.594640

**Authors:** Maria Loconsole, Silvia Benavides-Varela, Lucia Regolin

## Abstract

If you hear the non-words ‘Kiki’ and ‘Bouba’, you may be more likely to associate them with a spiky and a round object, respectively, rather than the opposite. This is a case of sound-symbolism, known as the Bouba-Kiki effect. Studies on four-months infants suggest that this effect might constitute a predisposed perceptual mechanism. However, these studies suffered from the impossibility of ruling out a fast experience-driven origin of the effect resulting from infants’ speed of learning, their sensitivity to environmental regularities, and the large number of sound-symbolic associations to which they are precociously exposed when interacting with adults. To better describe its ontogeny and fill in this gap, we tested the Bouba-Kiki effect in domestic chicks (*Gallus gallus*). Being a precocial species, chicks can be tested on the very early days of life, allowing for a virtually total control of their experience before test. Three-day-old chicks (n=42) first learned to circumnavigate a panel to obtain a food reward. Then, they were presented with two identical panels, one depicting a spiky shape, and one depicting a round shape, while hearing either the sound ‘Bouba’ or ‘Kiki’. We recorded which panel chicks chose with either sound, in a total of 24 trials. Chicks preferred the panel with the spiky shape when hearing the ‘Kiki’ sound, and that with the round shape when hearing the ‘Bouba’ sound. Results from naïve baby chicks hint at a predisposed mechanism for matching the two dimensions of shape and sound that may be widespread across species.

## Introduction

In his pioneering work in 1947, Wolfgang Kölher described the spontaneous tendency of adult humans to associate a round shape with the non-word ‘Maluma’ and the spiky shape with the non-word ‘Takete’ (1). This connection between a meaningless arbitrary sound and a likewise meaningless geometrical shape was interpreted as a form of sound-symbolism, namely a spontaneous crossmodal association between a sound (‘Takete’/’Maluma’) and a shape (spiky/round). Several studies replicated this effect, which, since the 2001 study by Ramachandran and Hubbard, was renamed the Bouba-Kiki effect (2), i.e., the non-words ‘Bouba’ and ‘Kiki’ were matched to the round and the spiky shape, respectively. Even though the Bouba-Kiki effect appears to be a strong and stable effect in our species, it is yet to know how the spontaneous tendency to match shapes and sounds originates. Some authors proposed that these associations could emerge due to experience conveyed by early exposure to multisensory information and word sounds (3), or by orthography: people might consider the sounds [b] and [o] rounder than the sounds [k] and [i] due to the shape of the letters (4, 5). Others postulated that the ability to connect linguistic labels that resemble the form of their referents is an experience-independent and spontaneous ability in our species (2, 6). According to the latter view, these form-to-meaning correspondences would constitute a predisposed perceptual mechanism at the basis of language acquisition, facilitating vocabulary construction and communication in infancy (7, 8). The strongest evidence in favour of this view comes from cross-linguistic and language evolution studies (9), as well as research in preverbal infants. The first attested the Bouba-Kiki effect in cultures with different linguistic systems and orthographic appearance of letters (10, 11). The latter showed that, before language production, infants (4 to 7 months old) already exhibit the Bouba-Kiki effect (12, 13). However, studies on infants are not conclusive, as they could not completely rule out a fast experience-driven origin of the effect resulting from infants’ speed of learning (14, 15), their high sensitivity to environmental statistical regularities (16), and the large number of symbolic associations of sounds to which they are exposed when interacting with adults (17). Although sound symbolism has never been directly investigated in non-human animals, comparative research on different species provided solid evidence of other instances of crossmodal correspondences, including associations between pitch and luminance (chimpanzees (18)), pitch and size (chimpanzees (19), dogs (20, 21) and tortoises (22)), and space and luminance (domestic chicken (23)). Taken together, these studies hint at a shared and evolutionary analogous mechanism to perceive and represent physical properties of objects according to common rules. This mechanism could confer an advantage to the individuals, allowing them to make accurate predictions about co-occurrences in the environment, reducing uncertainty, and allowing the formation of coherent and meaningful representations of objects and events, such as facilitating certain associations (e.g., small objects and high pitch sounds, or small objects and elevated spatial positions) while hindering those that are less frequent (24, 25). Sound symbolism, namely, the Bouba-Kiki effect, may constitute one of these shared crossmodal representations. Direct evidence of sound-symbolism in an animal model is crucial to clarify whether it belongs to a set of common predispositions that facilitate the organism-environment interaction and as such constitute one of the building blocks of knowledge or, rather, it is a uniquely human system of associations that results from language-related experiences.

Our study aims at describing both the ontogeny and the phylogeny of sound-symbolism within the comparative approach, testing the Bouba-Kiki effect in naïve newborn domestic chickens (*Gallus gallus*). Baby chicks represent an optimal model to this aim, as they are known to share analogous predisposed cognitive and perceptual mechanisms to human infants, such as intuitive physical reasoning (26, 27), sensitivity to animacy (28, 29) and to biological motion (30, 31). Three-day-old chicks were also tested for a case of crossmodal correspondence, that is, space-luminance association (23), performing similarly to adult humans (32) and showing a spontaneous tendency to match dark stimuli with the left hemispace and bright stimuli with the right hemispace. Additionally, results from research on baby chicks well added on studies on mammalian species, suggesting that crossmodal associations may reflect a similar prenatal organization and development of the involved neural mechanisms (19, 24, 25). Instances of a spontaneous crossmodal association (i.e., visual-tactile) have recently been identified at the onset of life, in chicks tested soon after hatching, suggesting that the brain may be spontaneously organised to allow such associations (33).

Here we trained naïve baby chicks to circumnavigate a panel to obtain a food reward. Then, we presented chicks with two identical panels, one depicting a spiky shape, and one depicting a round shape, while either the sound ‘Bouba’ or ‘Kiki’ was played in the background. We hypothesised that, if chicks rely on sound-symbolic associations similar to humans, they will prefer the panel with the spiky shape when hearing the ‘Kiki’ sound, and that with the round shape when hearing the ‘Bouba’ sound. Studying a precocial animal will allow full control of the experience prior to testing, overcoming one of the most debated limitations in infant studies. This approach allows to pinpoint the developmental origin of sound symbolism and clearly address the role of experience, if any, on its emergence. Additionally, evidence from a bird species can provide relevant insights on the phylogeny of this phenomenon, considering the wide phylogenetic distance between mammals and birds (with the last common ancestors being dated between 300 and 320 million years ago (34, 35)).

## Results

The best-fitting model for our data (i.e., the one having the lowest AIC value) was the one including the background sound as the only relevant predictor. This indicates that while chicks’ performance can be at least partially explained by the background sound (i.e., ‘Bouba’, or ‘Kiki’), there is no relevant effect of their sex (male, or female), the testing trial (i.e., performance remains stable across the 24 trials), the position (left, or right) of the stimuli, and their interactions. Post-hoc analysis on the selected model revealed that chicks significantly prefer the round shape when hearing the sound ‘Bouba’ (P(spiky) = 0.34, SE = 0.02, z = -7.01, p < 0.0001), and significantly prefer the spiky shape when hearing the sound ‘Kiki’ (P(spiky) = 0.56, SE = 0.02, z = 2.67, p = 0.01) (**Fig. 1**).

**Fig. 1:**
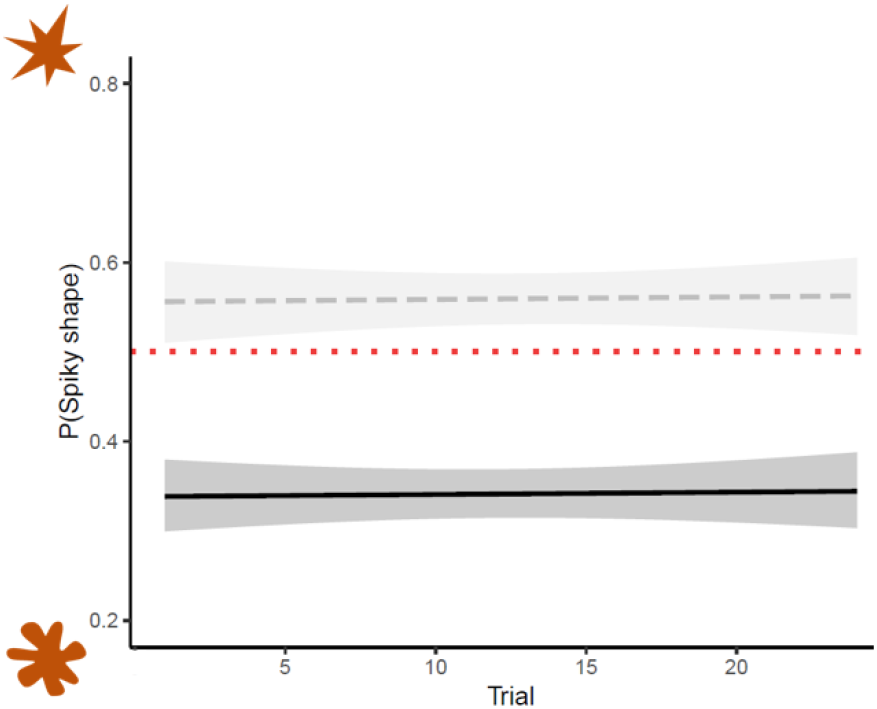
Results. The y-axis depicts the probability that an animal approaches the spiky shape. The x-axis shows the testing trials (from 1 to 24). The dashed grey line refers to the trials in which chicks were presented with the ‘Kiki’ sound; the continuous black line refers to the trials in which chicks were presented with the ‘Bouba’ sound. The shaded areas represent 95% confidence intervals. The dotted red line represents chance level (P = 0.5). Overall, chicks consistently chose significantly more often the panel with the spiky shape when hearing ‘Kiki’, and that with the round shape when hearing ‘Bouba’.

## Discussion

The present study tested baby naïve domestic chickens for an example of sound-symbolism, namely the Bouba-Kiki effect, aiming at shedding light on its ontogenetic origin. In fact, even if this is a strong and well attested effect in our species, in both adults (2) and infants (12, 13), there is still an active debate on how it originates. We reported a spontaneous tendency to match a round and a spiky shape to the ‘Bouba’ and ‘Kiki’ sounds, respectively, in three-day-old chicks. Our results fit in the current debate suggesting that sound-symbolism belongs to a set of predisposed associations built into different species. Our data well support the existence of an early-available mechanism as we hatched the eggs in the lab, where we exerted a virtually full control of chicks’ experience, overcoming one of the biggest limitations from studies on human infants. Even though our subjects had never experienced the sound-symbolic matching prior to test, they still spontaneously associated the two dimensions of shape and sound. This indicates that sound-symbolic associations (at least in the case of the Bouba-Kiki effect) may take place from the earliest stages of life, without the need of the subject directly experiencing the crossmodal matching. Similar results were obtained for other instances of crossmodal associations in this species, such as visual-spatial (23) and visual-tactile (33), pointing toward a predisposed general mechanism for crossmodal associations. Evidence from several non-human species, including chimpanzees (18), monkeys (36), dogs (20, 21), and tortoises (22), suggest that such a mechanism may be shared across different taxa, and possibly reflect an evolutionary old organising principle of the brain (18, 22). A possible advantage that comes from a predisposition for crossmodal associations is for naïve animals to readily respond to environmental statistical regularities, as several crossmodal associations mirror natural co-occurrences. For instance, in the case of the pitch-size association, there is a biological constraint for which smaller animals would produce higher sounds in pitch with respect to larger animals (due to the shape and size of their vocal tract): this allows for a reliable prediction of the expected size of a non-visible animal by simply hearing their call (19, 24, 37). For what concerns sound-symbolism, in humans a predisposition for matching a sound to a referent is thought to have a pivotal role in language emergence and vocabulary formation (7, 38). An endowed predisposition for associating speech sounds with their referents would help infants in creating a lexical representation of their surroundings and selectively focusing on referents in a complex environment, thus reducing computational loading (2, 38). Interestingly, however, previous studies showed that spontaneous crossmodal associations (space-luminance in chicks (23) and pitch-size in tortoises (22)) fade over repeated trials, eventually reaching chance level. This was interpreted considering that, to serve their scope, crossmodal priors must be flexible and update with experience, hence fading after repeated experiences of unrewarding trials. We did not observe this same pattern in our experiment. However, due to differences in the employed methodology it is difficult to address the origin of this difference. One possibility could be that by including refresh trials (rewarded) prior to each test block (unrewarded), as well as a time break between each block, we reinstated chicks’ attention and motivation toward the testing stimuli (39), thus preserving the initial association. Alternatively, it is possible that sound-symbolic associations are deeply rooted and thus less subject to habituation or extinction. Further studies should be devoted to disentangling between these two possibilities.

In light of our results, we suggest that a predisposition to sound-symbolic associations may support different communicative systems across species, and as such it is not unique to human language. In fact, there is growing evidence showing that non-human animals do possess complex systems of communications that share some basic properties with human language, including sound-to-referent mapping. For instance, chimpanzees produce different grunts to indicate different types of food (40); vervet monkeys emit distinct calls to signal the presence of different predators (snake, eagle, and leopard), each one eliciting an appropriate response in the other monkeys (respectively: inspecting the ground, hiding in a bush or under a tree, climbing up a tree) (41, 42); Japanese tits were shown to produce alarm calls to specifically signal the presence of a snake, resulting in the flock reacting more promptly to snake-like stimuli (and not to other predators) (43); and domestic chickens possess a wide repertoire of calls (44), emitting specific calls for different predators(45, 46). We suggest that, similarly to other instances of crossmodal associations, sound-symbolism also constitutes a predisposed set of associations that supports the first interactions that an organism has with their environment. In humans, this could represent a building block that prompts language development. In other species, it could have an analogous role, supporting the further development of a refined system for communication, that allows, for example, mentally representing and referring to distinct elements that are not in the immediate proximity (7, 8).

## Conclusions

In our study, we showed that baby domestic chickens (*Gallus gallus*) respond to sound-symbolic associations similar to humans, preferentially matching the sounds ‘Bouba’ and ‘Kiki’ to a round and a spiky shape, respectively. These results pave the way toward a clear and univocal description of the phenomenon of sound-symbolism by tackling both its ontogeny and its phylogeny at the same time. Evidence from a precocial species allows overcoming one crucial limitation of studies on infants, allowing a virtually full control of the animal’s experience prior to test. This makes it possible to pinpoint the developmental origin of sound-symbolism and, according to our results, places it at the earliest stages of life, possibly hinting at a predisposed experience-independent mechanism. Crucially, direct evidence in an animal model suggests that, rather than being a culturally learned phenomenon unique to humans, sound-symbolism may belong to a set of predisposed associations built into different species.

Even though this constitutes a first important step in settling the debate on the origin of sound-symbolism, further studies should be dedicated to better understanding the role of experience on the development of sound-to-referent mapping. The presence of similar mechanisms in chicks and human infants may be of particular interest for future work devoted at casting some light on the ultimate function of this ability and its role in the development of more sophisticated cognitive mechanisms.

## Materials and Methods

### Subjects and rearing conditions

The experiments complied with all applicable national and European laws concerning the use of animals in research and were examined and approved by the ethical committee of the University of Padua (Organismo Preposto al Benessere Animale - O.P.B.A.), N. Prot 795/2, N. Project 35/2024.

Fertilized eggs were obtained bi-monthly from a local hatchery and incubated (FIEM, MG 70/100 FAMILY) in the laboratory at controlled temperature (37.5° C) and humidity (50%). Upon hatching chicks were feather sexed and housed in same-sex social groups of three individuals in standard metal cages (28 × 32 × 40 cm). Cages were enlightened by 36 W fluorescent lamps placed 15 cm above the cage from 7am to 7pm and followed by 2/3h blocks of dark/light alternation from 7pm to 7am. At all times chicks had free access to water and food in the cage. Training and testing took place on day 3 of life. One hour before the beginning of the experimental procedures, chicks were food deprived, while water remained always available. At the end of the experimental procedures, chicks were donated to local farmers. We tested a total of 42 subjects, 17 of which were females.

### Training

The experimental arena consisted of a plastic triangle (76cm side x 32cm height, **Fig. 2**) similar to that employed in previous studies on crossmodal correspondences in chicks of the same age (23, 47). The aim of the training was to familiarise the chicks with the arena and teach them to circumnavigate an opaque panel (12cm base x 15cm height x 3.5cm thick) made of white cardboard to obtain a palatable food reward, i.e., half a mealworm (*Tenebrio molitor*). The panel depicted an ambiguous shape made of both spiky and round edges. At first the chick was presented with a single panel in the centre of the arena (**Fig. 2A**). Once the chick had learnt to promptly circumnavigate the panel when let free in the arena, it was exposed to two panels, one on the left and one on the right, one had the training shape printed on it (this was the rewarded panel) and the other was blank (**Fig. 2B**). The position of the rewarded panel was counterbalanced across subsequent training trials so that it never appeared on the same side more than twice consecutively. Once the chick completed 6 out of 8 consecutive correct trials (retrieving the food behind the panel depicting the ambiguous shape) it entered the test phase. On average, training lasted between 10 and 15 minutes per chick, requiring approximately 10 to 15 trials to each subject. Importantly, before test chicks were never exposed to the test shapes, nor they heard any distinctive sound (other than the calls of the other chicks in the rearing room) therefore it was very unlikely that they had reasons for associating any test shape to a sound.

**Fig. 2:**
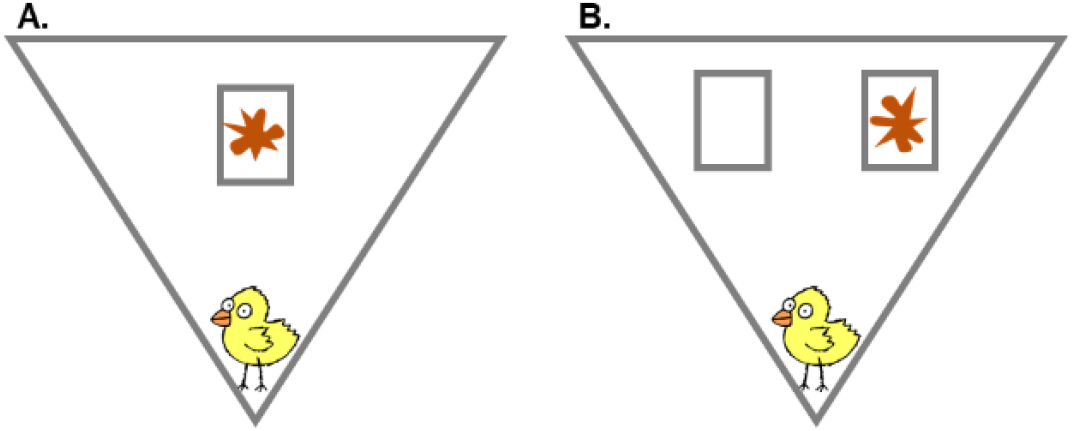
Training. **A**. The chick learned to circumnavigate a central panel depicting an ambiguous shape featuring both spiky and round edges to retrieve a palatable food. **B**. Thereafter the chick presented with two panels learned to circumnavigate the one depicting the ambiguous shape. The position (left/right) of the baited panel was counterbalanced between training trials. To pass the training phase, the chick had to correctly respond in 6 out of 8 consecutive trials.

### Test

The test arena is identical to the one used for training. Chicks were presented with both panels, which now depicted a round and spiky shape, respectively (**Fig. 3**). Each chick underwent four blocks of six trials each, for a total of 24 trials. During each trial, either the non-word ‘Bouba’, or ‘Kiki’ were played in the background via a small speaker (SONY SRSXB01, 8 x 5 x 5 cm) placed centrally behind the arena. The ‘Bouba’ and ‘Kiki’ sounds, as well as the round and spiky shapes were the same employed in a previous study on humans that tested for the Bouba-Kiki effect across different cultures and writing systems (11). There was no need to modify the auditory stimuli, as humans and chicks respond to similar auditory frequencies (48, 49). Similarly, chicks have a fine visual system (50, 51), for which they will be able to process the visual stimuli used for human studies without any foreseeable impediment. The colour of the shapes was however changed from black to orange, as chicks are more responsive, and more likely to pay attention to such colour (52). Before starting each trial, the chick was temporarily hold in the starting vertex behind a transparent partition. After hearing two repetitions of either the ‘Bouba’ or ‘Kiki’ sound (approximately three seconds) the chick was let free to move in the arena and approach one of the two panels, while the background sound kept playing. No reward was provided in any of the test trials. After the beginning of the trial, after lifting the transparent partition to release the chick in the arena, the experimenter remained away from the arena to a defined spot out of the chick’s sight, observing its behaviour from a camera on top of the arena. Once the trial was completed (i.e., the chick had circumnavigated one of the panels) the experimenter approached again the arena to gently move the chick to a transport box while preparing the setup for the subsequent trial. After completing a block of six trials the chick was returned to the cage to rest for approximately one hour (with water ad libitum but no food available). Then, the chick received a refresh session made of four trials with both screens, one of which depicted the ambiguous stimulus used at training. This re-training was needed to prevent loss of motivation during unrewarded test trials, correct choices were in fact rewarded with half a mealworm. At test the position of the two shapes and the sound played were pseudo-randomized across trials, so that the same condition never occurred twice consecutively.

**Fig. 3:**
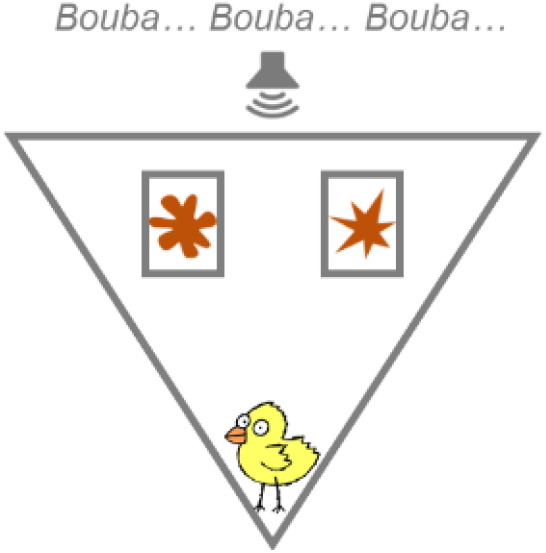
Test. An example of a testing trial (‘Bouba’ condition). The chick could see both a round and a spiky shape simultaneously presented each on one of the two panels, while a background sound repeated the non-word ‘Bouba’. If chicks possess predisposed sound-symbolic associations they might prefer to approach the shape congruent to the background sound, even if they had never experienced these stimuli before, neither individually nor together.

A choice was considered when the chick had circumnavigated a panel with its head and at least 2/3 of its body (23, 47). Once a choice had been made, the chick was gently removed from the arena without allowing it to inspect the panel not chosen, and a new trial started after a few seconds. The entire test was video recorded to allow offline scoring of the behaviour, the experimental response being the panel inspected by the chick on each trial.

### Data analysis

Raw data generated during the study are available as supplementary material (**S1**). Statistical analysis were run using R 4.2.1(53). We performed multiple nested linear mixed effect models with a binomial error structure (R package: lme4 (54)). The dependent variable was the shape depicted on the panel circumnavigated by the chick (0 = round, 1 = spiky). The independent variables were the background sound played during each trial (‘Bouba’ or ‘Kiki’), the trial number (from 1 to 24), chicks’ sex (male, or female), the position of the round stimulus (left, or right), and all the interactions between these factors. Individual chicks were included in the model as the random effect. We then ran an Akaike information criterion (AIC) based model selection to choose the best fitting model. We run a post-hoc analysis with Tukey correction on the selected model (R package: emmeans (55)) to determine the direction of effect of the predictors.

## Acknowledgments

We wish to thank Francesca Simion for discussing the original idea of this study, and Tommaso Feraco for his suggestions on the project draft.

Maria Loconsole’s work is funded by the European Union – NextGenerationEU and by the University of Padua under the 2023 STARS Grants@Unipd programme (project *CROSS - Comparative Research Of Sound Symbolism)*. Silvia Benavides-Varela and Lucia Regolin are founded by the Italian Ministry of Education and Research through the Research Project of National Relevance (PRIN) – 2022 Prot. 2022TYX52H.

## Author Contributions

**M.L**.: Conceptualization, Methodology, Validation, Formal analysis, Investigation, Writing – Original Draft, Visualization, Funding acquisition. **S.B.V**.: Conceptualization, Writing – Review & Editing, Funding acquisition. **L.R**.: Conceptualization, Methodology, Resources, Writing – Review & Editing, Funding acquisition.

## References

1. W. Köhler, Gestalt psychology; an introduction to new concepts in modern psychology, Rev. ed (Liveright, 1947).

2. V. S. Ramachandran, E. M. Hubbard, Synaesthesia--a window into perception, thought and language. Journal of Consciousness Studies 8, 3–34 (2001).

3. D. J. Lewkowicz, A. A. Ghazanfar, The emergence of multisensory systems through perceptual narrowing. Trends in Cognitive Sciences 13, 470–478 (2009).

4. G. Lockwood, M. Dingemanse, Iconicity in the lab: a review of behavioral, developmental, and neuroimaging research into sound-symbolism. Frontiers in Psychology 6 (2015).

5. A. Nielsen, D. Rendall, The sound of round: Evaluating the sound-symbolic role of consonants in the classic Takete-Maluma phenomenon. Canadian Journal of Experimental Psychology / Revue canadienne de psychologie expérimentale 65, 115–124 (2011).

6. R. Bottini, M. Barilari, O. Collignon, Sound symbolism in sighted and blind. The role of vision and orthography in sound-shape correspondences. Cognition 185, 62–70 (2019).

7. M. Dingemanse, D. E. Blasi, G. Lupyan, M. H. Christiansen, P. Monaghan, Arbitrariness, iconicity, and systematicity in language. Trends in Cognitive Sciences 19, 603–615 (2015).

8. P. Perniss, G. Vigliocco, The bridge of iconicity: from a world of experience to the experience of language. Philosophical Transactions of the Royal Society B: Biological Sciences 369, 20140179 (2014).

9. D. E. Blasi, S. Wichmann, H. Hammarström, P. F. Stadler, M. H. Christiansen, Sound–meaning association biases evidenced across thousands of languages. Proceedings of the National Academy of Sciences 113, 10818–10823 (2016).

10. A. J. Bremner, et al., “Bouba” and “Kiki” in Namibia? A remote culture make similar shape–sound matches, but different shape–taste matches to Westerners. Cognition 126, 165–172 (2013).

11. A. Ćwiek, et al., The bouba/kiki effect is robust across cultures and writing systems. Philosophical Transactions of the Royal Society B: Biological Sciences 377, 20200390 (2021).

12. M. Fort, A. Weiss, A. Martin, S. Peperkamp, Looking for the bouba-kiki effect in prelexical infants in (2013).

13. O. Ozturk, M. Krehm, A. Vouloumanos, Sound symbolism in infancy: evidence for sound-shape cross-modal correspondences in 4-month-olds. J Exp Child Psychol 114, 173–186 (2013).

14. J. S. Horst, L. K. Samuelson, Fast Mapping but Poor Retention by 24-Month-Old Infants. Infancy 13, 128–157 (2008).

15. K. Wanrooij, P. Boersma, T. Van Zuijen, Fast phonetic learning occurs already in 2-to-3-month old infants: an ERP study. Frontiers in Psychology 5 (2014).

16. J. R. Saffran, N. Z. Kirkham, Infant Statistical Learning. Annual Review of Psychology 69, 181–203 (2018).

17. C. V. Parise, F. Pavani, Evidence of sound symbolism in simple vocalizations. Exp Brain Res 214, 373–380 (2011).

18. V. U. Ludwig, I. Adachi, T. Matsuzawa, Visuoauditory mappings between high luminance and high pitch are shared by chimpanzees (Pan troglodytes) and humans. Proceedings of the National Academy of Sciences 108, 20661–20665 (2011).

19. A. K. Kalan, R. Mundry, C. Boesch, Wild chimpanzees modify food call structure with respect to tree size for a particular fruit species. Animal Behaviour 101, 1–9 (2015).

20. A. T. Korzeniowska, J. Simner, H. Root-Gutteridge, D. Reby, High-pitch sounds small for domestic dogs: abstract crossmodal correspondences between auditory pitch and visual size. Royal Society Open Science 9, 211647 (2022).

21. A. T. Korzeniowska, H. Root-Gutteridge, J. Simner, D. Reby, Audio–visual crossmodal correspondences in domestic dogs (Canis familiaris). Biology Letters 15, 20190564 (2019).

22. M. Loconsole, G. Stancher, E. Versace, Crossmodal association between visual and acoustic cues in a tortoise (Testudo hermanni). Biology Letters 19, 20230265 (2023).

23. M. Loconsole, M. S. Pasculli, L. Regolin, Space-luminance crossmodal correspondences in domestic chicks. Vision Research 188, 26–31 (2021).

24. V. F. Ratcliffe, A. M. Taylor, D. Reby, Cross-Modal Correspondences in Non-human Mammal Communication. Multisens Res 29, 49–91 (2016).

25. L. Dj, G. Aa, The emergence of multisensory systems through perceptual narrowing. Trends in cognitive sciences 13 (2009).

26. R. Baillargeon, J. Li, W. Ng, S. Yuan, “An account of infants’ physical reasoning” in Learning and the Infant Mind, (Oxford University Press, 2009), pp. 66–116.

27. C. Chiandetti, G. Vallortigara, Intuitive physical reasoning about occluded objects by inexperienced chicks. Proc Biol Sci 278, 2621–2627 (2011).

28. E. Di Giorgio, M. Lunghi, G. Vallortigara, F. Simion, Newborns’ sensitivity to speed changes as a building block for animacy perception. Sci Rep 11, 542 (2021).

29. O. Rosa-Salva, M. Grassi, E. Lorenzi, L. Regolin, G. Vallortigara, Spontaneous preference for visual cues of animacy in naïve domestic chicks: The case of speed changes. Cognition 157, 49–60 (2016).

30. F. Simion, L. Regolin, H. Bulf, A predisposition for biological motion in the newborn baby. Proceedings of the National Academy of Sciences 105, 809–813 (2008).

31. G. Vallortigara, L. Regolin, F. Marconato, Visually Inexperienced Chicks Exhibit Spontaneous Preference for Biological Motion Patterns. PLOS Biology 3, e208 (2005).

32. A. Fumarola, et al., Automatic spatial association for luminance. Atten Percept Psychophys 76, 759–765 (2014).

33. E. Versace, L. Freeland, S. Wang, M. G. Emmerson, First-sight recognition of touched objects shows that chicks can solve the Molyneux’s problem. [Preprint] (2022). Available at: https://www.biorxiv.org/content/10.1101/2022.08.18.504388v1 x[Accessed 29 January 2023].

34. N. J. Emery, N. S. Clayton, Evolution of the avian brain and intelligence. Current Biology 15, R946–R950 (2005).

35. E. D. Jarvis, et al., Avian brains and a new understanding of vertebrate brain evolution. Nat Rev Neurosci 6, 151–159 (2005).

36. A. A. Ghazanfar, J. G. Neuhoff, N. K. Logothetis, Auditory looming perception in rhesus monkeys. Proceedings of the National Academy of Sciences 99, 15755–15757 (2002).

37. A. M. Taylor, D. Reby, K. McComb, Human listeners attend to size information in domestic dog growls. J Acoust Soc Am 123, 2903–2909 (2008).

38. M. Imai, S. Kita, The sound symbolism bootstrapping hypothesis for language acquisition and language evolution. Philosophical Transactions of the Royal Society B: Biological Sciences 369, 20130298 (2014).

39. R. Rugani, M. Loconsole, L. Regolin, A strategy to improve arithmetical performance in four day-old domestic chicks (Gallus gallus). Sci Rep 7, 13900 (2017).

40. K. E. Slocombe, K. Zuberbühler, Functionally Referential Communication in a Chimpanzee. Current Biology 15, 1779–1784 (2005).

41. R. M. Seyfarth, D. L. Cheney, P. Marler, Vervet monkey alarm calls: Semantic communication in a free-ranging primate. Animal Behaviour 28, 1070–1094 (1980).

42. R. M. Seyfarth, D. L. Cheney, P. Marler, Monkey Responses to Three Different Alarm Calls: Evidence of Predator Classification and Semantic Communication. Science 210, 801–803 (1980).

43. T. N. Suzuki, Animal linguistics: Exploring referentiality and compositionality in bird calls. Ecological Research 36, 221–231 (2021).

44. N. E. Collias, The Vocal Repertoire of the Red Junglefowl: A Spectrographic Classification and the Code of Communication. The Condor 89, 510–524 (1987).

45. C. S. Evans, L. Evans, P. Marler, On the meaning of alarm calls: functional reference in an avian vocal system. Animal Behaviour 46, 23–38 (1993).

46. C. S. Evans, J. M. Macedonia, P. Marler, Effects of apparent size and speed on the response of chickens, Gallus gallus, to computer-generated simulations of aerial predators. Animal Behaviour 46, 1–11 (1993).

47. M. Loconsole, A. Gasparini, L. Regolin, Pitch–Luminance Crossmodal Correspondence in the Baby Chick: An Investigation on Predisposed and Learned Processes. Vision 6, 24 (2022).

48. D. Purves, et al., The Audible Spectrum. Neuroscience. 2nd edition (2001).

49. E. M. Hill, G. Koay, R. S. Heffner, H. E. Heffner, Audiogram of the chicken (Gallus gallus domesticus) from 2 Hz to 9 kHz. J Comp Physiol A Neuroethol Sens Neural Behav Physiol 200, 863–870 (2014).

50. H. M. Johnson, Visual pattern-discrimination in the vertebrates -II. Comparative visual acuity in the dog, the monkey and the chick. Journal of Animal Behavior 4, 340–361 (1914).

51. R. Over, D. Moore, Spatial acuity of the chicken. Brain Research 211, 424–426 (1981).

52. A. d Ham, D. Osorio, Colour preferences and colour vision in poultry chicks. Proceedings of the Royal Society B: Biological Sciences 274, 1941–1948 (2007).

53. R Core Team, R: A Language and Environment for Statistical Computing. (2022).

54. D. Bates, M. Mächler, B. Bolker, S. Walker, Fitting Linear Mixed-Effects Models Using lme4. Journal of Statistical Software 67, 1–48 (2015).

55. R. V. Lenth, et al., emmeans: Estimated Marginal Means, aka Least-Squares Means. (2023). Deposited 23 June 2023.

